# Cortical-wide impairment of “The Glymphatic system” after focal brain injury

**DOI:** 10.1101/2022.10.05.510560

**Authors:** M. Azuma, H. Lee, K. Shinzaki, R. Yamane, M. Morita

## Abstract

In peripheral tissues, cells are maintained in the interstitial fluid that flows from capillaries to lymph system. However, the brain has no lymphatic capillaries, and the actual state of the interstitial fluid has long been unknown. Recently, a glymphatic system has been proposed in which part of the cerebrospinal fluid flowing on the surface of brain tissue enters the brain parenchyma via the peri-arterial space, becomes interstitial fluid, and then flows out again from the peri-venous space. Brain injury due to head contusion or stroke is thought to impair the intracerebral circulation and aggravate the extracellular environment, but the actual situation is unknown. Therefore, in this study, we examined the effects of focal brain tissue damage on intracerebral circulation using the light-injured mouse, an originally developed closed head injury model. In light-injured mice, the injury-making process does not affect intracerebral circulation because the cranium is maintained. However, this method has quantitative problems, so we developed a method to image cerebrospinal fluid and blood vessels from the surface of the cerebral cortex. After examining different injury sites and different time periods after injury, it was found that intracerebral circulation was reduced to the same extent on the ipsilateral and contralateral sides of the injury at one-week post-injury. This intracerebral circulatory deficit was still partially present at four-weeks post-injury. These results indicate that the intracerebral circulation is extensively impaired by local injury and neurodegenerative diseases.

## Introduction

The flow of interstitial fluid in the brain, which lacks capillary lymphatics, has long been unknown, but a Glymphatic system integrating cerebrospinal fluid and interstitial fluid has recently been proposed (Iliff et al., 2012). This system is based on the idea that cerebrospinal fluid produced in the ventricular choroid plexus flows through the subarachnoid space into the brain parenchyma from the spaces around the penetrating arteries to become cerebral interstitial fluid, then flows out from around the veins, and finally drains out of the brain through the cribriform plate (Kida et al., 1993; Norwood et al., 2019), subarachnoid flow (Pollay, 2010), and meningeal lymph vessels (Ahn et al., 2019; Louveau et al., 2015, 2018), etc. This process is accelerated by sleep and anesthesia. This flow is accelerated by sleep and anesthesia (Hablitz et al., 2019, 2020; Xie et al., 2013) and is dependent on AQP4 in astrocytes(Ishida et al., 2022; Simon et al., 2022, p. 4). It is also regarded as a mechanism to remove extracellular beta-amyloid, the causative agent of Alzheimer's disease (Da Mesquita et al., 2018; Iliff et al., 2014; Ishida et al., 2022; Peng et al., 2016; Reeves et al., 2020; Simon et al., 2022).

Focal brain injury associated with head contusion or cerebral infarction is expected to cause neurological and psychiatric disorders by inducing Glymphatic system failure and accelerating β-amyloid accumulation. In fact, it is widely accepted that brain injury is a risk for the development of Alzheimer's disease (Bolte et al., 2020; Iliff et al., 2014). It has already been reported that brain edema in the acute phase of brain injury compresses the flow pathways of the Glymphatic system and inhibits its flow (Mestre et al., 2020; Schain et al., 2017). It is thought that the Glymphatic system is also altered during the recovery period of brain injury due to the activation of glial cells, infiltration of immune cells, and production of extracellular matrix related to tissue repair (Mckee & Daneshvar, 2015; Monai et al., 2019), as well as changes in the distribution of AQP4, but the actual state of the Glymphatic system is unclear. Particularly, it is known that injury can alter neural activity not only in the proximity of the injury but also distally known as diaschisis, and it is expected that this is mediated by a failure of the Glymphatic system (Bolte et al., 2020; Iliff et al., 2014).

In the present study, we examined Glymphatic system dysfunction during the chronic phase of injury using our originally developed mouse model of closed head trauma, the photic injury mouse. Photo injury is a method of destroying a part of the cerebral cortex by irradiating intense light through a thin section of the cranium, and it is possible to create injuries in various brain regions without damaging the cranium, which constitutes a part of the perivascular space on the brain surface (Morita, 2014; Suzuki et al., 2012). We thought that this advantage would reveal how the circulation in the brain goes from failure to recovery, depending on the location of the brain injury and the time course after the brain injury. In conventional histological studies of the Glymphatic system, tracers such as fluorescently labeled dextran are injected into cerebrospinal fluid and analyzed based on the distribution of fluorescence in tissue sections. However, this method only quantifies cerebrospinal fluid distributed in some penetrating vessels in the tissue sections and cannot evaluate the wide range of Glymphatic system failure associated with brain injury. For this reason, in this study, we developed a method to comprehensively image blood vessels without sectioning brain tissue and used this method in our investigations. Specifically, the cerebral cortex was sectioned, flattened, and made transparent, and the distribution of blood vessels and cerebrospinal fluid from the surface to the depth was imaged using confocal microscopy. If the effects of brain injury on intracerebral circulation can be clarified through this study, it is expected to provide not only clinically important findings but also a new perspective on the mechanism of intracerebral circulation, which is currently the subject of active debate.

## Materials and Methods

### Mice

All animal experiments were approved by the institutional animal care and use committee of Kobe University (Permission number: xx-xx) and performed in compliance with the Kobe University Animal Experimentation Regulations. C57BL/6J mice between two and six months old were used.

### Photo injury

Thinned cranial window was created on right hemisphere after anesthetizing mice by the intramuscular injection of ketamine (100mg/kg, Daiiichi-sankyo, Tokyo, Japan) and xylazine (10 mg/kg, Bayer Yakuhin, Osaka, Japan) and incising scalp, by scraping skull of 1 mm in diameter to 20 μm thick by drill and dental knife. The location of cranial window was 1.5 mm posterior to bregma and 4 mm lateral from midline for photo injury on middle cerebral artery or 3 mm for that on somatosensory cortex. Photo injury was induced by irradiating cerebral cortex through the cranial window and microscope objective (UMPlanFI 20 X Olympus, Tokyo, Japan) with a halogen lamp at 26 mW for 2.5min. Then, scalp was sutured, returned to animal facility after recovery from anesthesia and maintained for the periods designated.

### Labeling of cerebrospinal fluid and blood vessel

Mice were anesthetized with the ketamine and xylazine, fixed in a stereotaxic device (NARISHIGE, Tokyo, Japan) with head tiled down at 50°, maintained at 36°C by heating pad. Glass capillaries (G150F-3, Warner Instruments, Hamden) were pulled by a micropipette puller (Model P-87, SUTTER INSTRUMENT, Novato, USA) and their tips were mechanically crushed by approximately 2 mm. These capillaries were filled with 5 μL of saline containing TAMRA-Ovalbumin (10mg/mL), connected to injector (IMS-10 15009, Narishige), inserted 2 mm between skull and spine to reach cistern magna by a micromanipulator (MS-10 15009, Narishige). The tracer was injected at 2.5-3.3 μL/min. Mice were euthanized by cardiac arrest due to injecting 500 μL of 1.24 M KCl into heart chamber and glass capillaries were gently removed one minute after the termination of breath. Mice were fixed by the cardiac perfusion of PBS and 4% paraformaldehyde warmed at 37°C, descending aorta was clamped and then 500 μl warmed PBS containing Cy5-Ovalbumin (18.8mg/ml) and gelatin (0.1g/ml, Nippi, Tokyo, Japan) were perfused at 13.5 mL/min.

### Tissue clearance

Cerebral hemispheres of fixed brains were dissected, flattened by compressed between two glass plates of 2 mm distance and cleared by SeeDB (Ke et al., 2013), which was submerging in 20%, 40%, and 60% (wt/vol) fructose solutions containing 0.5% α-thioglycerol2 for 4 to 8 hours, 80% and 100% fructose solutions for 12 hours and SeeDB solution(80.2% fructose) for 24 hours respectively with gentle agination at room temperature.

### Imaging

Fluorescence images were obtained by an epifluorescence microscope (IX70, Olympus) equipped with a cooled-CCD camera (ORCA-R2, Hamamatsu Photonics, Hamamatsu, Japan) or confocal laser scan microscopy (Fluoview1000, Olympus). Z-stack confocal images were obtained every 20 μm of approximately 700 μm thick flattened cerebral hemisphere, by optimizing laser power and PMT sensitivity. A Fiji plugin, Stitching was used for stitching of confocal images. Bright filed images were obtained by a microscope (CK40, Olympus, Tokyo, Japan) equipped with a camera (ILCE-QX, Sony, Tokyo, Japan).

### Extraction of vascular structure from fluorescence images and subsequent quantification of CSF

Surface vessels were extracted as structures larger than 500 –1000 pixels (approximately 3,000 – 6,000 μm^2^) in upper half images of the Z-stack, after binarization depending on thresholds determined by experimenter. Penetrating vessels were extracted as structures that span more than three layers (60μm) and connected to surface vessels after enhancement of boundary by high pass filter using Fourier transform, subtraction of surface vessels and size base noise reduction with criteria that penetrating vessels are between 16 pixels (100μm^2^), which is typical size of capillary (Carare et al., 2008) and 500 –1000 pixels (approximately 3,000 – 6,000 μm^2^), which is below the surface vessel size described above.

### Immunohistochemistry

Mice were fixed by the cardiac perfusion of ice-cold PBS containing heparin (1000mL, Wako, Osaka, Japan) and 4 % paraformaldehyde PBS after anesthesia by intraperitoneal injection of pentobarbital (75mg/kg). Brains were extracted and coronally sliced at 100μm thickness by a vibratome (DOSAKA, Osaka, Japan). Slices were subjected to a series of treatment at room temperature; 50mM NH_4_Cl for 10 min, two PBS wash for 10 min, 0.4% Triton X-100 for two hours, three PBS wash for 5 min, and blocking with PBS containing 5% normal goat serum and 0.1% Triton X-100 (blocking solution) for one hour, then incubated with primary antibody diluted to blocking solution for overnight at 4°C. Slices were returned to room temperature and subjected to three PBS wash for 10 min, incubation with secondary antibody in PBS for two hours, three PBS wash for 10 min, and mounting in glycerol or Gelvatol (Harlow and Lane, 2006) containing 0.1% p-phenylenediamine. Primary antibodies were rabbit anti-AQP4 (1:250;Vector Laboratories), guinea-pig anti-GFAP (1:250;Vector Laboratories), and mouse anti-S100ß (1:250;Vector Laboratories). Secondary antibodies were purchased from Abcam and Jackson ImmunoResearch.

### Cervical lymph node collection

After cisternal magna injection of Evans Blue (20%) or TAMRA-Ova, mice were fixed, and lymph node were harvested.

### Fluorescent protein preparations

2.5 mL of 10 mg/mL Cy5 or TAMRA in DMSO were gently added to 2 mL of 20 mg/mL Ovalbumin in 0.2M NaHCO3 (pH8.4), slowly mixed for one hour and then purified by PD-10 column (**).

### Reagents

If not specified, reagents purchased from nacalai tesque (Kyoto, Japan).

## Results

### Fluorescence images of CSF and blood vessel (BV)

CSF and blood vessel (BV) were fluorescently labeled by injecting TAMRA-conjugated ovalbumin (TAMRA-Ova) into cisterna magna and vascular cast with Cy5-labelled ovalbumin (Cy5-Ova). Low magnification images demonstrate TAMRA-labelled CSF (CSF / TAMRA-Ova) and Cy5-labelled BV (BV / Cy5-Ova) (Fig.1A). The fluorescence of CSF and BV was preserved after flattening of cerebral cortex and tissue clearing (Fig.1B). Confocal images were obtained every 20 μm depth and surface (Surface) or maximum projection (Z project) images were stitched to illustrate CSF and BV of whole cerebral cortex (Fig.1C). CSF along penetrating arteriole and venule were compared, because the line of literature on glymphatic system suggests the accumulation of fluorescence tracer of CSF in arteriole prior to venule. Representative arteriole (A) and venule (V) in the highlighted area in Fig 1C / Z project were circled at surface, where A and V were connected to surface vessel, and depths from 0 μm, at which penetrating vessels were identified to 600 μm (Fig. 1D), and their fluorescent images were aligned every 20μm in Fig.1E. These results suggests that the CSF fluorescence venule was limited within 100 μm, while that of arteriole reached 600 μm. This point was statistically confirmed by comparing the fluorescence intensity of CSF normalized by that of BV (F_CSF_/F_BV_) between arteriole and venule at each depth (Fig 1F). In this paper, the difference between arteriole and venule were not further discussed, because the majority of penetrating vessels were hardly assigned to arteriole or venule. Instead, we developed an algorithm for extracting the structure of penetrating vessels and comprehensively analyzed CSF along them. Our algorithm extracted 4685 penetrating vessels from seven hemispheres of seven mice, and their F_CSF_/F_BV_ were shown after sorting by maximum depth of each vessel (Fig. 1G). The split violin plot of F_CSF_/F_BV_ at 0 μm and at 200 μm (Fig 1G inset) indicates two distinguished components at 0 μm, which may reflect arteriole and venule, while a single component at 200 μm, which may reflect arteriole. This non-normal distribution suggest the requirement of non-parametric statistical test for comparing F_CSF_/F_BV_ between mouse groups.

**Fig.1.**
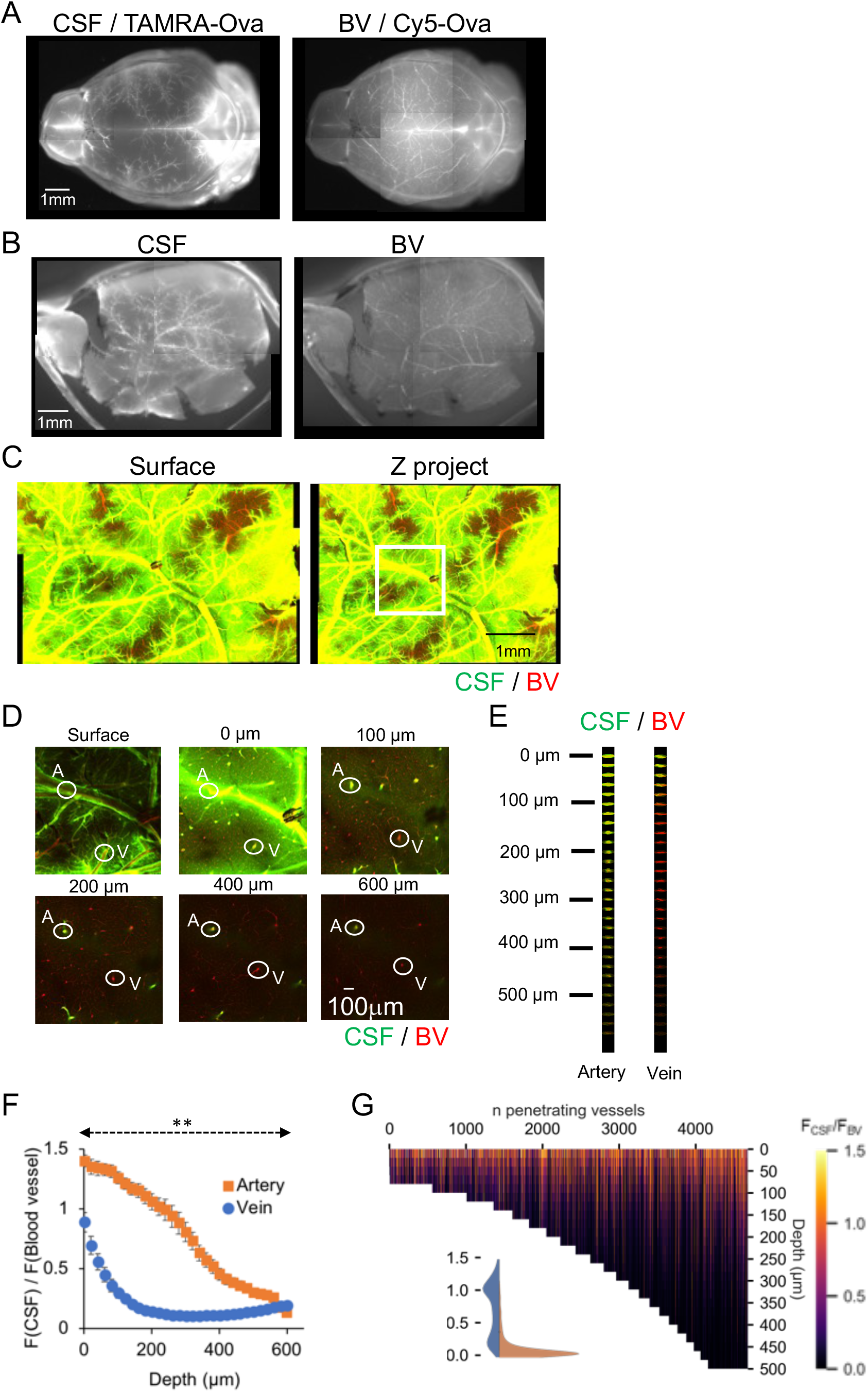
Fluorescence labelling of cerebral vasculature and CSF. A. Whole brain fluorescence images of blood vessel (BV) and CSF. TAMRA- or Cy5-labelled ovalbumin were injected in cisternal magna or left ventricle respectively. B. Fluorescence images of BV and CSF in a cerebral hemisphere after flattening and tissue clearing. C. Stitched confocal images of BV (red) and CSF (green) in surface (left) and whole depth (right) of hemisphere. Confocal images obtained by using a 10x objective as described in Materials and Methods, and subjected tiling. Representative image including surface vessel were used for “surface”, while images were used after maximum projection for “whole depth”. D. Representative images at surface, 100 μm, 200 μm, 400 μm, 600 μm in depth. Typical penetrating artery (A) and vein (V) were circled. Fluorescence images of CSF and BV of the artery and vein in (D) were aligned with depth. F. CSF distributions along arteries and veins in all depth. CSF was quantified as F_CSF_/F_BV_, which is the fluorescence intensity of CSF normalized by that of BV. Means±SE (10 arteries and 10 veins from six mice), **p<0.01. G. Heatmap of F_CSF_/F_BV_ along individual vessels sorted by maximum depth. Violin plot of F_CSF_/F_BV_ at surface (blue) and 200μm in depth (orange). n = 4685 form 7 hemispheres.

### Broad impairment of the glymphatic system after photo injury on mid-cerebral artery

In order to investigate the influence of focal brain injury on the glymphatic system, the distribution of fluorescently labelled CSF was examined seven days after photo injury, when edema is largely dissolved, and tissue recovery starts (Morita, 2014).Initially, injury was created on middle cerebral artery (Fig. 2A, arrow on MCA/7dpi) as the most severe case including reduced cerebral blood perfusion. As a result, CSF bilaterally decreased in broad area of cerebral cortex in comparison with a normal mice shown Fig 1. Surface CSF (Fig. 2B) was largely decreased and limited along major arteries both in ipsilateral (MCA / 7dpi / ipsi) and contralateral (MCA / 7dpi / contra) hemispheres. The similar decrease and limitation of CSF was found in z-projection image (Fig.2C), suggesting the reduction of CSF in the branches of surface arteries. Since the BV fluorescence was lost only in lesion core (Fig.2C arrow), the branches of surface arteries possess intact blood while the loss of CSF. Comprehensive analysis of surface BV and CSF indicates that the brain injury significantly reduced CSF in both hemispheres, and the reduction was significantly larger in ipsilateral hemisphere (Fig 2D). This was also the case in the comprehensive quantification of CSF along penetrating vessels presented as heatmap (Fig. 2E) and following statistical analysis (Fig.2F). Overall, focal brain injury on MCA impaired the glymphatic system in broad area of cerebral cortex over both hemispheres, while the impairment was statistically more severe in ipsilateral. The impaired CSF in the contralateral side implicates that the influence of focal brain injury on the glymphatic system is not solely mediated by vasculature.

**Fig.2.**
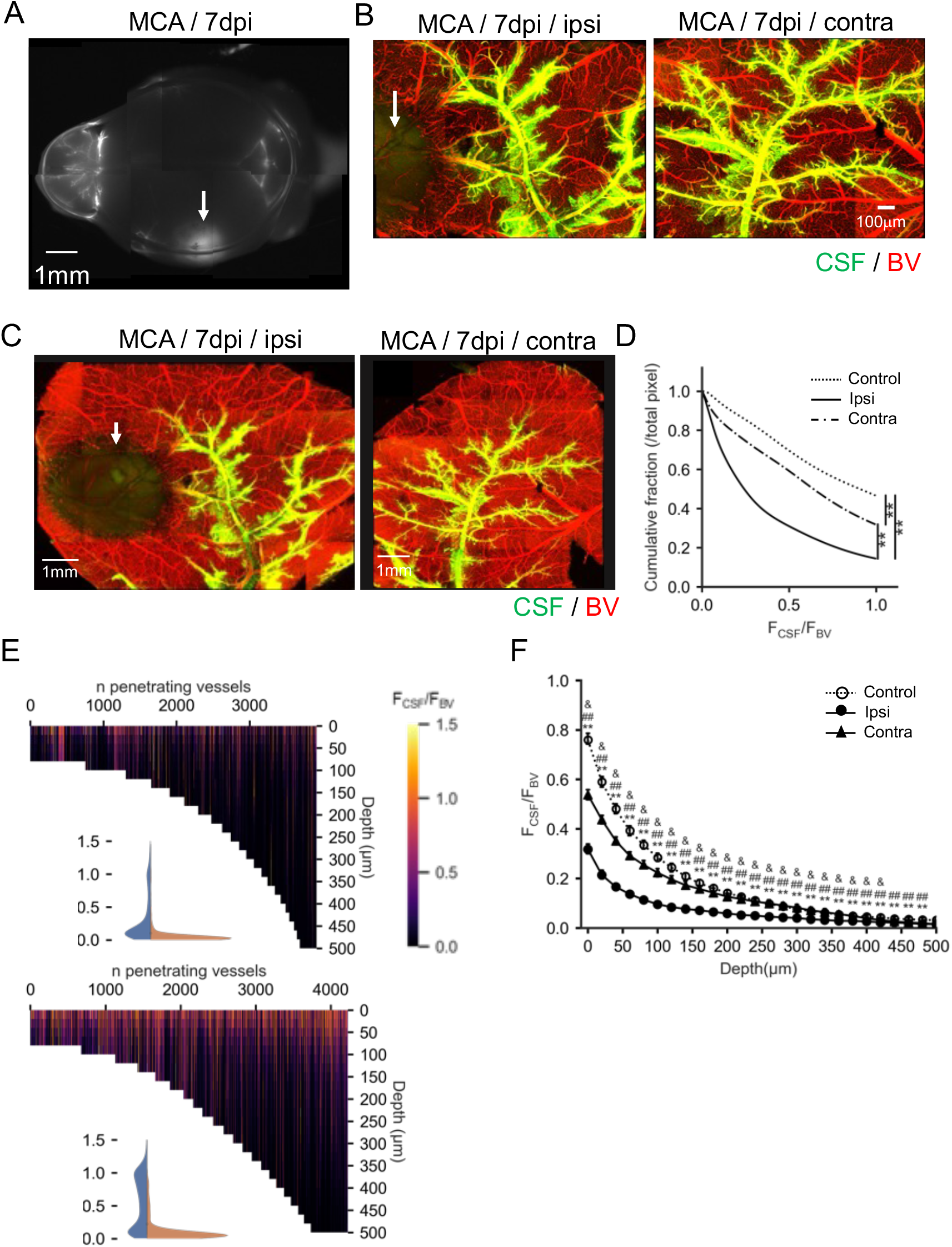
Broad impairment of glymphatic system following photo injury on middle cerebral artery. Seven days after photo injury on middle cerebral artery (MCA/7dpi). A. Whole brain fluorescence image of CSF. Lesion is marked with an arrow. B. Surface images of BV and CSF in ipsilateral (MCA / 7dpi / ipsi) and contralateral (MCA/ 7dpi / contra) hemisphere. Lesion is marked with an arrow. C. Z stack images of BV and CSF in Maximum intensity projection for whole depth of ipsilateral (left) and contralateral (right) side of the hemispheres. Lesion site marked with an arrow. D. Distributions of CSF along surface vessels, shown as reverse cumulative curve of normalized pixel fractions vs F_CSF_/F_BV_. Kolmogorov-Smirnov test, **p<0.001. E. Entire vessels’ heatmap, ipsilateral hemisphere (up) and contralateral hemisphere (down), n = 3918 form 6 hemispheres from ipsilateral and n = 4226, 6 hemispheres from contralateral side. F. Distributions of CSF along penetrating vessels. Means±SE, DSCF post hoc test, *: control vs ipsi, #: control vs cont, &: ipsi vs cont, *p<0.05, **p<0.001

### Broad impairment of the glymphatic system after photo injury on somatosensory cortex

The influence of focal brain injury with limited alterations of cerebral blood circulation on the glymphatic system was examined seven days after photo injury created by avoiding major arteries on somatosensory cortex (Fig. 3A, SSC / 7dpi, arrow). The size of lesion core was smaller after photo injury on SSC in comparison with that on MCA, but broad reduction of CSF was similarly found. Surface CSF was largely decreased and limited along major arteries both in ipsilateral (SSC / 7dpi / ipsi) and contralateral (SSC / 7dpi / contra) hemispheres (Fig. 3B), and similar decreases and limitation of CSF was found in z -projection image (Fig. 3C). Comprehensive analysis of surface BV and CSF showed significant bilateral reduction of CSF, and the extent of reduction was significantly larger in ipsilateral hemisphere (Fig 3D). This was also the case in the comprehensive quantification of CSF along penetrating vessels presented as heatmap (Fig. 3E) and following statistical analysis (Fig. 3F).

**Fig.3.**
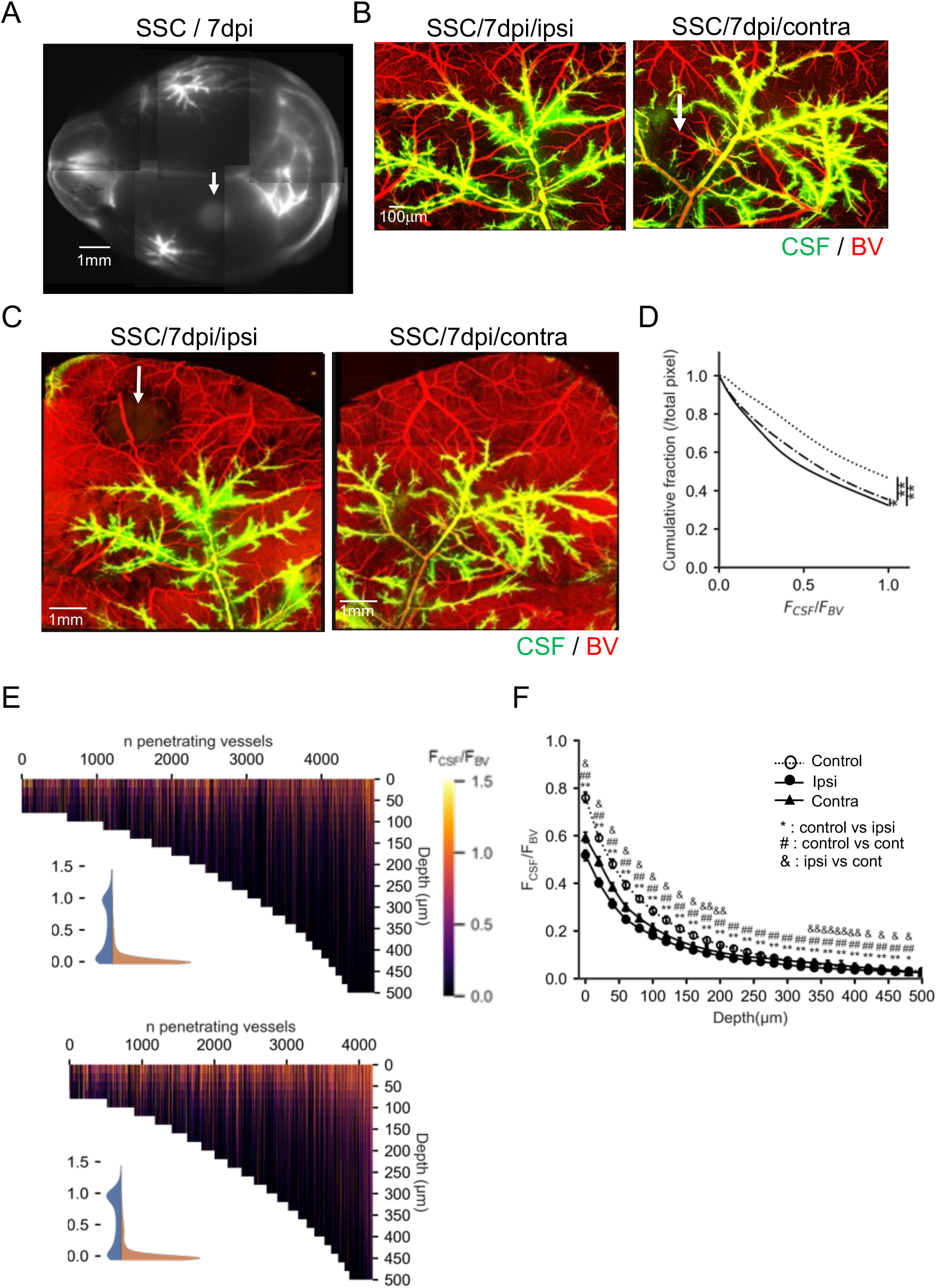
Broad impairment of glymphatic system following photo injury on somatosensory cortex. Seven days after photo injury on somatosensory cortex (SSC/7dpi). A. Whole brain fluorescence image of CSF. Lesion is marked with an arrow. B. Surface images of BV and CSF in ipsilateral (SSC/ 7dpi / ipsi) and contralateral (SSC/ 7dpi / contra) hemisphere. Lesion is marked with an arrow. C. Z stack images of BV and CSF in Maximum intensity projection for whole depth of ipsilateral (left) and contralateral (right) side of the hemispheres. Lesion site marked with an arrow. D. Distributions of CSF along surface vessels, shown as reverse cumulative curve of normalized pixel fractions vs F_CSF_/F_BV_. Kolmogorov-Smirnov test, **p<0.001. E. Entire vessels’ heatmap, ipsilateral hemisphere (up) and contralateral hemisphere (down), n = 4709 form 7 hemispheres from ipsilateral and n = 4188, 5 hemispheres from contralateral side. F. Distributions of CSF along penetrating vessels. Means±SE, DSCF post hoc test, *: control vs ipsi, #: control vs contra, &: ipsi vs contra, *p<0.05, **p<0.001

### Recovery of glymphatic system after 28 days after photo injury on somatosensory cortex

The influence of recovery from focal brain injury on the glymphatic system was examined on 28 days after photo injury. As a results, lesion was still visible (Fig.4A, arrow on SSC / 28dpi), but CSF recovered to the similar lever as in normal mice. Both surface and Z-projection images also showed robust recovery of CSF (Fig. 4B and C), and lesion without BV fluorescence largely shrunk (Fig 4.C, SSC/28dpi/ipsi, arrow). Comprehensive analysis of surface BV and CSF still showed significant reduction in ipsilateral hemisphere, but the CSF of contralateral side was indistinguishable from normal mice (Fig. 4D). CSF was even significantly increased after recovery in both hemisphere in comparison with control as in the comprehensive quantification of CSF along penetrating vessels presented as heatmap (Fig. 4E) and following statistical analysis (Fig. 4F). Overall, the reduced CSF after photo injury on SSC was mostly recovered in surface vessels and even increased along penetrating vessels by 28 days.

**Fig.4.**
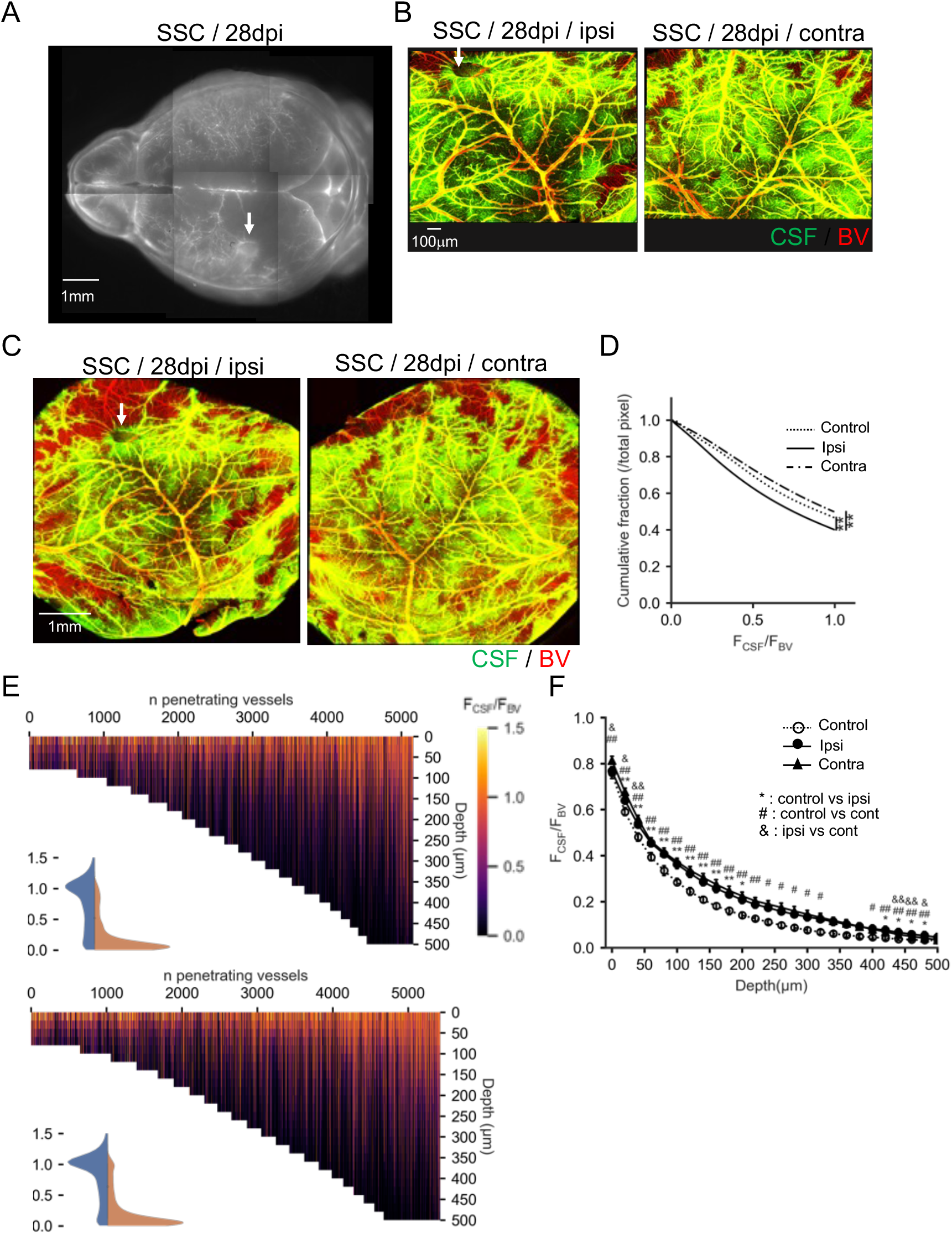
Recovery of glymphatic system after photo injury on somatosensory cortex. Twenty-eight days after photo injury on somatosensory cortex (SSC/28dpi). A. Whole brain fluorescence image of CSF. Lesion is marked with an arrow. B. Surface images of BV and CSF in ipsilateral (SSC/ 28dpi / ipsi) and contralateral (SSC/ 28dpi / contra) hemisphere. Lesion is marked with an arrow. C. Z stack images of BV and CSF in Maximum intensity projection for whole depth of ipsilateral (left) and contralateral (right) side of the hemispheres. Lesion is marked with an arrow. D. Distributions of CSF along surface vessels, shown as reverse cumulative curve of normalized pixel fractions vs F_CSF_/F_BV_. Kolmogorov-Smirnov test, **p<0.001. E. Entire vessels’ heatmap, ipsilateral hemisphere (up) and contralateral hemisphere (down), n = 5160 form 7 hemispheres from ipsilateral and n = 5426, 6 hemispheres from contralateral side. F. Distributions of CSF along penetrating vessels. Means±SE, DSCF post hoc test, *: control vs ipsi, #: control vs contra, &: ipsi vs contra, *p<0.05, **p<0.001

### AQP4 expression after the brain injury

Since AQP4 knockout mice shows impaired glymphatic system as after photo injury, the reduction or altered localization of AQP4 was examined by immunohistochemistry. Cerebral cortexes of normal mice (Control) or ipsilateral hemisphere of mice 7 days after photo injury on SSC (SSC / 7dpi / ipsi) were stained byantibodies for an astrocyte marker S100ß, reactive astrocyte marker GFAP or AQP4 (Fig. 5A). As results, astrocyte activation was reflected in the reduction of S100b and the increase of GFAP after injury. AQP4 immunoreactivity was broadly increased after injury, while the vascular localization was preserved. As no reduction of AQP4 immunoreactivity was found, the hypothesis that the reduction of vascular AQP4 underlies the broad reduction of CSF after injury was excluded.

**Fig. 5.**
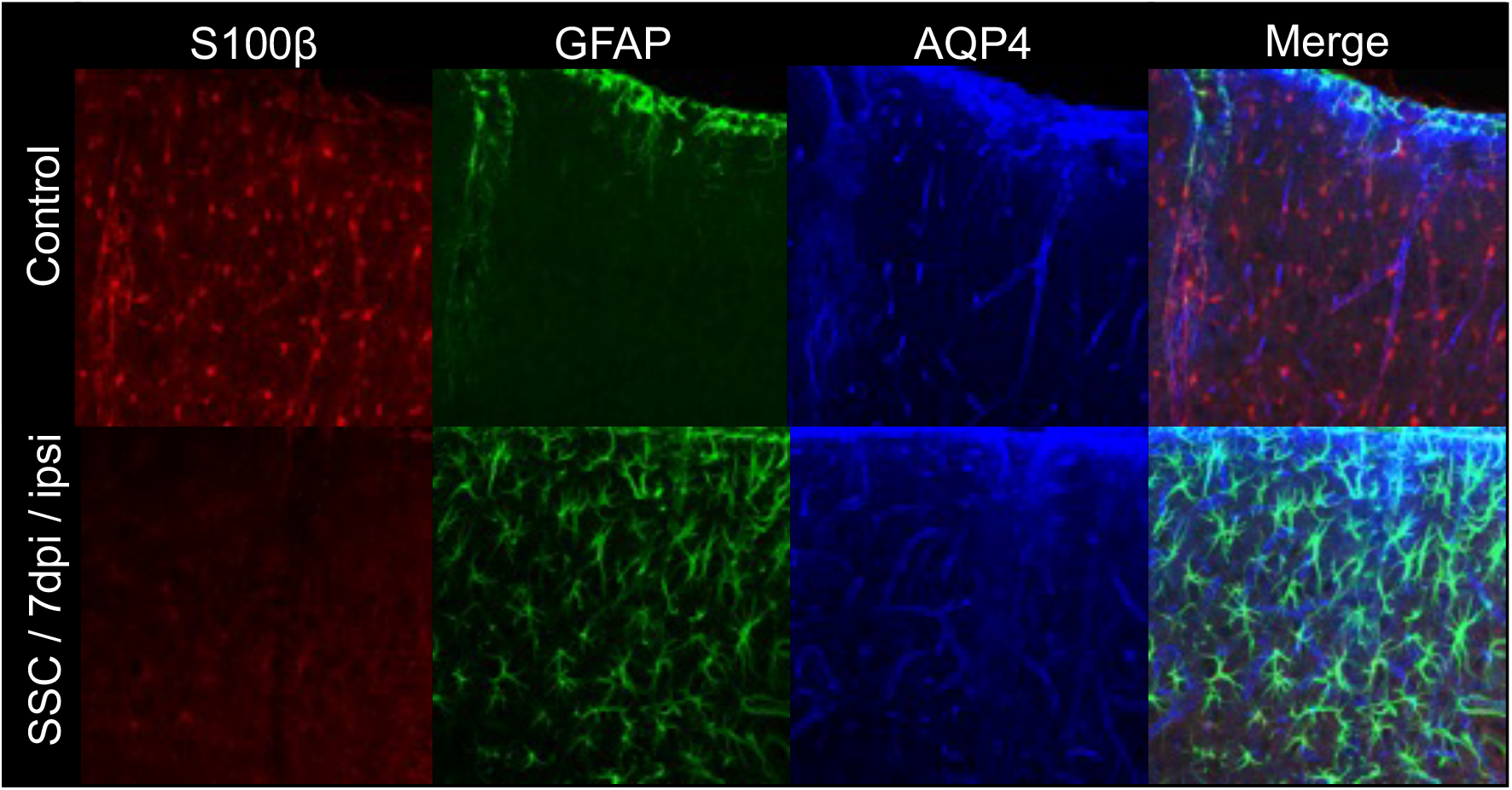
Astrocyte activation and AQP4 expression. Immunohistological analysis of astrocyte markers and AQP4 in normal mice (Control) and, mice at seven days after photo injury on SSC (SSC / 7dpi / ipsi). S100β (red), GFAP (green) and AQP4 (blue).

**Fig. 6.**
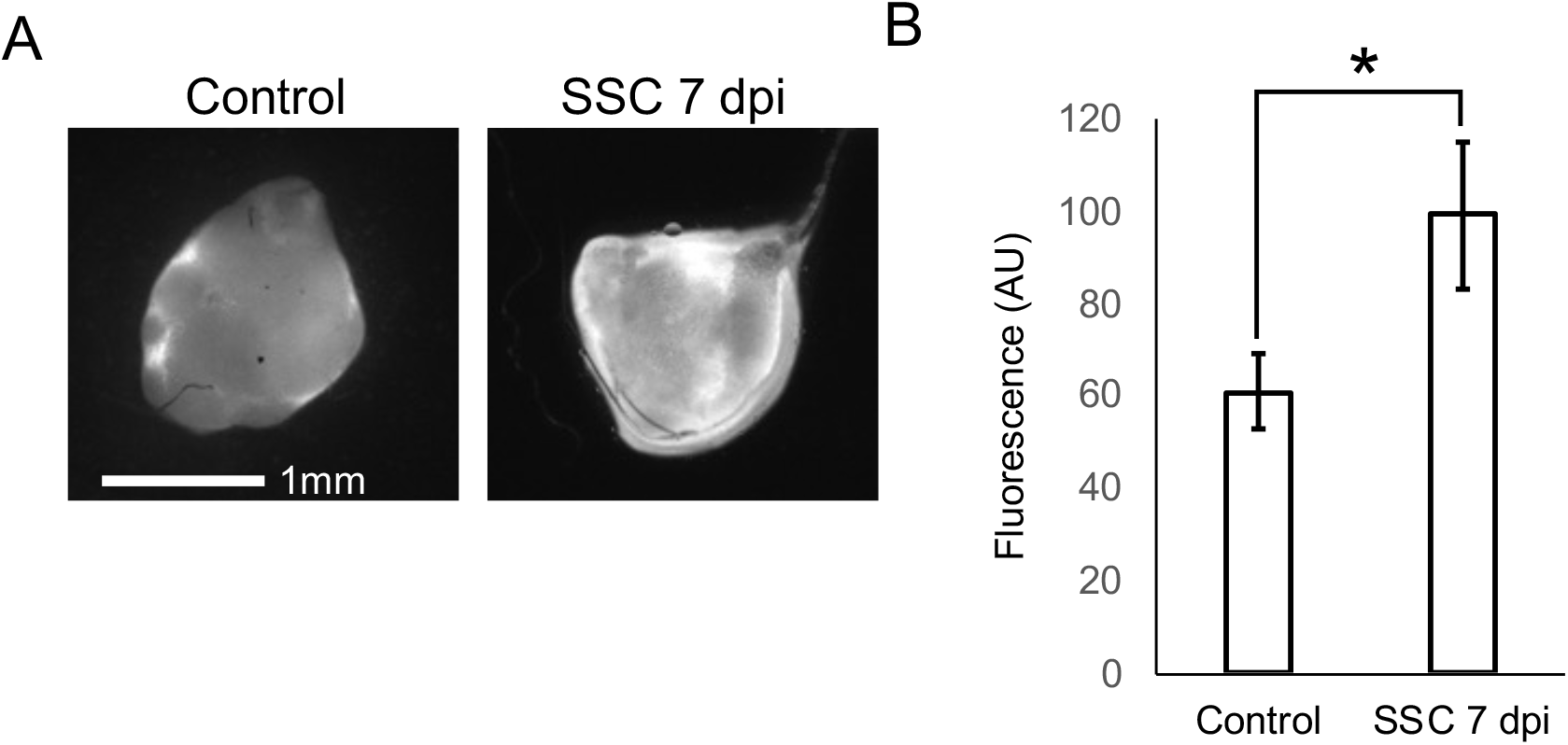
The upregulation lymphatic flow after photo injury. Lymphatic flow was evaluated by the accumulation of intracisternally-injected tracers in cervical lymph node. A. Fluorescence image of cervical lymph nodes after intracisternal injection of TAMRA-labelled ovalbumin (TAMRA-Ova). B. Quantitative analysis of fluorescence in cervical lymph node. Means±SE, t-test, n=3

### Lymphatic flow after the brain injury

We also hypothesized that brain injury impaired glymphatic system by suppressing the efflux of CSF into lymphatic system, and measured CSF drainage into lymphatic flow. Fluorescence of cervical lymph node after intracisternal injected fluorescence tracer was more intense in mice seven days after photo injury on somatosensory cortex (SSC / 7 dpi) than in normal mice (Control), indicating the upregulation of lymphatic drainage of CSF after brain injury. These results suggest two possibilities. One is that the upregulated drainage through meningeal lymphatic system reduced the CSF flow from major artery to brain parenchyma. This possibility is consistent with the substantial CSF in major surface arteries, while significant reduction in penetrating vessels. Another one is that the upregulation of drainage located along basilar artery, including cervical nerve, and subsequent reduction of CSF flows upward to cerebral cortex.

## Discussion

In this study, we established a method to quantitatively analyze cerebrospinal fluid distribution in individual vessels by double-labeling cerebrospinal fluid and vessels in the perivascular space using fluorescent tracers. In this method, penetrating arteries were labeled with cerebrospinal fluid to a greater depth than penetrating veins, confirming cerebrospinal fluid dynamics consistent with the Glymphatic system. Further examination of cerebrospinal fluid dynamics in light-injured mice revealed that cerebrospinal fluid distribution around the penetrating artery was significantly reduced in a wide range of brain regions, even at 1 week after injury, when brain edema had subsided. At 4 weeks of injury, when the condition had stabilized, cerebrospinal fluid distribution on the brain surface returned to a nearly normal state, but some insufficiency remained in the penetrating artery.

Intracerebral circulation has been studied by analyzing the distribution of tracers injected into the cerebrospinal fluid. A number of histological analyses of brain sections using fluorescent substances as tracers (Yang et al., 2013) and MRI using gadolinium and other substances as tracers (Gaberel et al., 2014) have already been reported. In the case of brain sections, the distribution of low molecular weight tracers soaked into the brain parenchyma or the distribution of relatively high molecular weight tracers that do not directly soak into brain tissue in the perivascular space has been used as an indicator. However, the low molecular weight probe soaks are unevenly distributed in the vicinity of the tracer injection site, making it difficult to provide an indicator of cerebrospinal fluid dynamics in the brain as a whole. MRI can image the tracer over time in a living state, but the resolution is extremely low (several centimeters), making it difficult to perform experiments using small animals such as mice, and it is difficult to image the entire length of the perivascular space in the same brain slice. In addition, it is impossible to analyze the structure of blood vessels and other organs. Based on the above results, this study developed a method to image cerebrospinal fluid distributed throughout the entire length of the space around blood vessels without using brain sections. In particular, by double-labeling cerebrospinal fluid and cerebral blood vessels and taking their ratio, we corrected for image degradation and other problems in the deep cerebral cortex. In addition, by comprehensively imaging the entire cerebral cortex, surface and penetrating blood vessels were simultaneously analyzed. Furthermore, by quantifying the cerebrospinal fluid distributed in the detected penetrating vessels using an algorithm to extract the vessels, we were able to successfully conduct cerebrospinal fluid at the microscopic level in a wide area of the brain.

Injury on the middle cerebral artery reduced cerebrospinal fluid distributed in both surface and penetrating vessels in a broad region spanning the ipsilateral and contralateral sides. Injuries made over the somatosensory cortex, avoiding the major vessels, similarly reduced cerebrospinal fluid distributed over both the superficial and penetrating vessels, while injuries over the middle cerebral artery reduced cerebrospinal fluid distributed over both the superficial and penetrating vessels. Regardless of the site of injury, the insufficiency of intracerebral circulation was comparable ipsilaterally and contralaterally. Although extensive failure of the intracerebral circulation with injury over the middle cerebral artery, including contralateral, has already been reported (Iliff et al., 2014), the ipsilateral failure is more severe. Equally extensive astrocyte activation, including altered localization of AQP4, has also been identified, which has been claimed as the cause of the failure (Iliff et al., 2014). The localization changes of AQP4 in this study were limited to the area surrounding the injury, making it unlikely that the increased failure is related to astrocyte activation. The beating of cerebral blood vessels has been claimed as the driving force for cerebrospinal fluid circulation, and evidence of this has been shown for intracerebral circulatory insufficiency associated with internal carotid artery tuberculosis (Iliff et al., 2013). It is also possible that with brain injury, blood vessels are damaged and blood pressure is lowered throughout the brain, resulting in reduced beating and cerebrospinal fluid flow. In this study, brain injury did not affect the fluorescent labeling of cerebral blood vessels, but its effect on cerebral blood flow remains unknown to date. It is highly possible that localized cerebral vascular damage associated with photic injury could affect blood flow throughout the brain, and it is conceivable that reduced cerebral blood flow and other factors could cause widespread intracerebral circulatory failure.

It has long been believed that the brain lacks lymphatic vessels, but recently the presence of lymphatic vessels in the meninges on the brain surface has been reported (Aspelund et al., 2015), and furthermore, meningeal lymphatic insufficiency has been shown to cause cerebrospinal fluid dynamic insufficiency and Alzheimer's disease development (Da Mesquita et al., 2018). Therefore, the possibility that these meningeal lymphatic vessels are the final drainage sites of cerebrospinal fluid has received significant attention. In this study, we examined the drainage of cerebrospinal fluid into the lymphatic system following brain injury and found that cerebrospinal fluid migration to the lymph nodes was largely unaffected, even under conditions in which cerebrospinal fluid distribution in the brain was greatly reduced. It is unlikely that cerebrospinal fluid drainage from the meningeal lymphatics on the same brain surface would occur at the same level as in normal mice, despite the reduction of cerebrospinal fluid on the brain surface due to photodamage. Thus, the results of this study suggest that lymphatic vessels are present in areas of high distribution of labeled cerebrospinal fluid in light-injured mice, such as the base of the brain, and are connected to the cervical lymph nodes.

The widespread failure of cerebral circulation seen 1 week after light injury likely increases the risk of memory impairment, depression, and epilepsy, which are known to be related to brain injury, due to the accumulation of waste products and ionic environmental imbalances (Pierce et al., 1998; Salazar et al., 1985). Cerebrospinal fluid is involved in the removal of potassium ions and lactate, which are released extracellularly with neural activity (Ball et al., 2010; Johanson et al., 2008). Accumulation of potassium ions increases neuronal excitability, leading to the development of epilepsy. Acidification of brain tissue associated with lactic acid accumulation is known to cause neural hyperactivity and other adverse effects on brain function (Halim et al., 2008). Although these symptoms are temporary and often diminish with recovery from the injury, the development of epilepsy and depression should be avoided because of their impact on post-recovery life. If methods to improve circulation in the brain are developed, it is desirable to select and treat patients with these symptoms.

In the present study, we also found a failure of intracerebral circulation in the penetrating artery even 4 weeks after injury. This insufficiency likely promotes the accumulation of beta-amyloid and is a risk factor for Alzheimer's disease. Dobutamine, an adrenergic agonist, has been reported to increase cardiac output, thereby increasing cerebral vascular pulsation and accelerating cerebrospinal fluid movement. After four-weeks of injury, when cerebral blood vessels have recovered sufficiently, activation of cerebral circulation with such drugs may be effective in avoiding the risk of Alzheimer's disease, while monitoring the insufficiency of cerebral circulation by MRI. We hope that this study will lead to improved recovery from brain injury.

## Notes

### Competing Interest Statement

The authors have declared no competing interest.

